# SpatialZoomer: multi-scale feature analysis of spatial transcriptomics

**DOI:** 10.1101/2025.09.08.674870

**Authors:** Xinqi Li, Yuhan Fan, Yue Han, Wenbo Guo, Jingmin Huang, Nan Yan, Zeyu Chen, Yanhong Wu, Yuxin Miao, Lin Hou, Xuegong Zhang, Zeyu Chen, Jin Gu

**Affiliations:** BNRIST Bioinformatics Division, Department of Automation, Tsinghua University, Beijing, China; MOE Key Laboratory of Bioinformatics, Tsinghua University, Beijing, China; Department of Computer Science and Technology, Tsinghua University, Beijing, China; Department of Statistics and Data Science, Tsinghua University, Beijing, China; School of Life Sciences, Tsinghua University, Beijing, China; School of Basic Medical Sciences, Peking University Health Science Center

## Abstract

Single-cell resolution spatial transcriptomics (ST) provides a great opportunity to explore the complex cellular contents in different tissues. In this study, we introduce SpatialZoomer, a spectral graph-based method that applies a set of low-pass filters to efficiently extract spatial molecular features from ST data at multiple resolutions or scales, including the single cell scale, the niche scale with dozens of closely interacting cells, and the domain scale with spatially-organized cell contents. The corresponding “critical” scales can be automatically identified by partitioning a cross-scale similarity map. Results show that SpatialZoomer can identify disease-progression signals at specific scales in Alzheimer’s Disease, and spatial context dependent cell subtypes in tumor microenvironment. The extracted multi-scale features also uncovered spatially heterogeneous niches in cancer. SpatialZoomer also takes advantage of high computational efficiency and low hardware requirements.

## Introduction

Spatial transcriptomics (ST) is an emerging technology which can detect the spatially resolved molecular information at single-cell or sub-cellular resolution. Recent advances in imaging-based technologies, such as 10x Genomics Xenium^1^, Vizgen MERFISH^2^, and Nanostring CosMx SMI^3^, combine *in situ* mRNA hybridization with labeled probes and cell segmentation to simultaneously profile hundreds to thousands of genes across millions of cells. These advances open opportunities to investigate cells’ molecular and morphological features, as well as spatial organization patterns.

Different types of cells are highly organized at multiple scales as functional units in different tissues. Recent studies have demonstrated the importance of multi-scale analytical strategies for ST data^4^. For example, a cell and its closely interacting neighbors may form a niche, while thousands of cells together can form a spatial functional unit or domain, such as tertiary lymphoid structures (TLSs) in cancer^5^. It has been demonstrated that a recurrent five-layered organization can be observed in glioblastoma samples by defining intercellular spatial relationships across scales, including colocalization, adjacency, and regional composition^6^. Numerous computational methods have been developed to characterize ST data at specific spatial scales, excelling in tasks such as cell type identification from subcellular or mixed-cell-resolution data^7,8^, niche detection^9,10^, and domain segmentation^11-15^. Some recent multi-scale algorithms have been proposed that vary the neighborhood sizes of circles or bins to simultaneously detect structures ranging from subcellular features and individual cell types to broader domains across different spatial scales^16-18^. However, most existing methods remain focused on finding predefined patterns at specific scales, without fully leveraging the varying scale as an additional analytical axis. Moreover, scale selection is often based on empirical parameter tuning rather than automated or adaptive strategies.

To address these gaps, we present SpatialZoomer, a computational framework for multi-scale analysis of single-cell resolution ST data, with scales automatically selected in a data-driven manner. This approach draws inspiration from zoom lenses in telescopes: zooming out reveals global structures, such as mountain contours, while zooming in uncovers local details, such as trees on the mountain or birds perched on their branches. SpatialZoomer extracts multi-scale spatial features using a bank of heat kernel-based spectral graph filters. As low-pass filters, heat kernels attenuate high-frequency components in the spectral domain while integrating spatial neighborhood contexts into cellular omics profiles in the vertex domain, with the scale parameter controlling the neighborhood size. Furthermore, SpatialZoomer automatically determines critical scales using a dynamic programming strategy that partitions the cross-scale similarity map into blocks, from which “representative” and “tipping” scales are identified. SpatialZoomer thereby extracts features and uncovers spatial structures across these critical scales.

We demonstrate SpatialZoomer’s capability through three applications. First, as different scales capture different signals, SpatialZoomer can reveal biologically meaningful patterns from noisy ST data that emerge only at specific scales. In longitudinal Alzheimer’s Disease (AD) tissue sections, disease progression-related signals are scarcely detectable at small scales but become apparent at certain scales. Second, SpatialZoomer enhances cell subtype identification by incorporating broader spatial contexts with transcriptomic profiles as scale increases. In a lung cancer dataset with 377 genes, it separates alveolar and interstitial macrophage subtypes, effectively mitigating the limitations of low gene coverage. In an ovarian cancer sample with 5,101 genes, it further identifies two cancer-associated fibroblast (CAF) subtypes with distinct spatial distribution, neighboring cell composition, transcriptomic profiles, and cell morphology, which remain indistinguishable in the raw transcriptomic space. Third, SpatialZoomer provides systematic insights into complex tissue architecture by jointly analyzing multi-scale spatial features. In cervical cancer, it reveals a stress-proliferation-EMT tri-layer organization and spatially heterogeneous niches. Moreover, SpatialZoomer can process datasets with millions of cells on a desktop computer within a few hours, offering high scalability, computational efficiency, and low hardware requirements. A multi-scale demo is available at https://li-xinqi.github.io/SpatialZoomer/, and the tool is accessible at http://lifeome.net/software/spatialzoomer.

## Results

### Overview of SpatialZoomer

SpatialZoomer is designed to extract multi-scale spatial features from single-cell resolution spatial omics data using spectral graph signal processing. It first constructs a spatial neighbor graph based on the physical proximity of cells, with node signals defined as cell programs derived from omics profiles (Fig. 1a). In the spectral domain, graph frequency characterizes the rate of signal variation across nodes: high-frequency components reflect rapid and local fluctuations between neighboring cells, while low-frequency components capture slow and global variations over larger spatial ranges. To extract spatial features at different scales, SpatialZoomer employs the heat kernel, a low-pass filter that exponentially attenuates high-frequency components while preserving low-frequency components, thereby smoothing the signals. From the vertex-domain perspective, this process simulates signal propagation following heat diffusion along the graph, where larger scale parameters lead to broader propagation and enable each cell to incorporate spatial contexts from larger neighborhood. The scale parameter of the heat kernel controls the filtering strength or the neighborhood size: at scale zero, the signal reflects the original single-cell scale; as the scale increases, signals from closely neighbors are incorporated, revealing niche-scale structures, and at larger scales, broader domain-scale patterns. Notably, the scale parameter of the heat kernel aligns with the concept of spatial scale, representing biological patterns at varying neighborhood sizes. By applying a bank of heat kernels with different scale parameters, SpatialZoomer enables simultaneous extraction of multi-scale spatial features ranging from local details to global structures (Fig. 1a).

**Figure 1.**
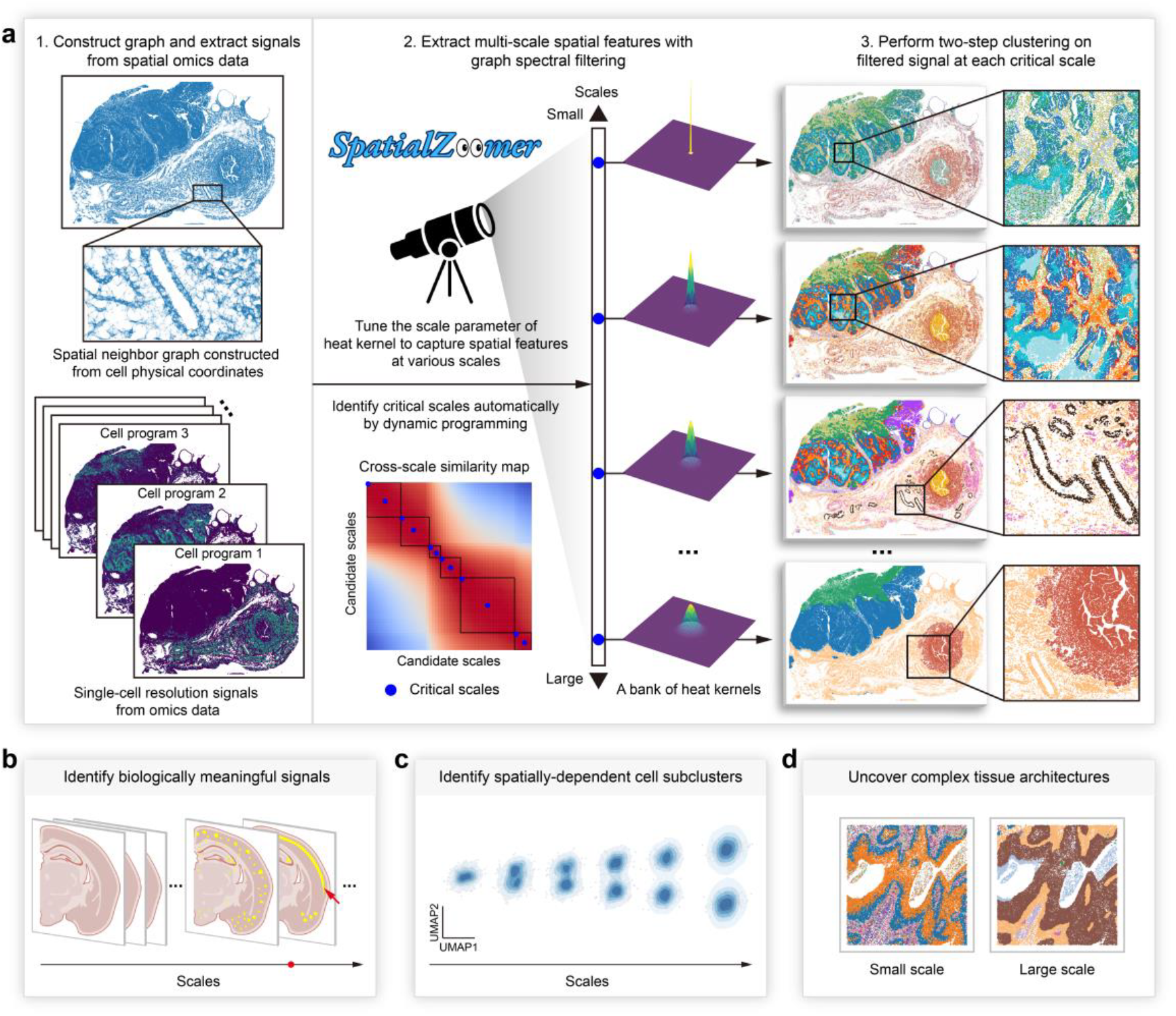
Schematic overview of SpatialZoomer. **a**, SpatialZoomer extracts multi-scale spatial features from single-cell resolution spatial omics data using spectral graph signal processing. Cell locations are employed to construct a spatial neighbor graph, while omics data are dimensionally reduced via Non-negative Matrix Factorization (NMF) to derive cell programs that serve as node signals. Tuning the heat kernel’s scale parameter extracts signals from neighborhoods of varying sizes, thereby capturing spatial features at different scales. A bank of spectral graph filters, constructed from heat kernels with different scale parameters, simultaneously extracts these multi-scale features. Critical scales are automatically identified from candidate scales by partitioning the cross-scale similarity map into blocks using dynamic programming. Two-step clustering is then performed on the transformed signals at these critical scales. The clustering results demonstrate SpatialZoomer’s application to an ovarian cancer sample profiled with the Xenium Prime platform, revealing distinct spatial structures at different critical scales. **b**-**d**, SpatialZoomer supports diverse downstream analyses.

Different spatial features emerge at different scales, analogous to how mountains, trees, and birds come into focus at varying focal lengths through a telescope. To automatically identify these critical scales, SpatialZoomer employs a dynamic programming (DP) strategy that partitions the cross-scale similarity map, which is constructed from transformed signals across ranked candidate scales, into contiguous diagonal blocks (Fig. 1a). The optimal number of blocks is also determined automatically. From each block, the start and center points are selected as critical scales, representing “tipping” and “representative” points, respectively.

Based on the transformed signals at these critical scales, SpatialZoomer applies a two-step clustering strategy (Fig. 1a). K-Means clustering is first performed to generate a large number of metacells, followed by the Leiden algorithm to assign final cluster labels (Fig. 1a). This strategy combines the computational efficiency of K-Means for large datasets with the adaptability of Leiden clustering for nonlinear manifold data. Additionally, it mitigates the spatial autocorrelation effects observed in graph based methods such as Leiden clustering and UMAP visualization of smoothed signals^19^. Moreover, the range of scale selection is not unlimited. At excessively large scales, spatial smoothing overwhelms omics heterogeneity, resulting in block-like clusters on the spatial slide and a tendency toward spherical patterns in UMAP space that lack biologically meaningful distinctions.

By quantitatively assessing the relative contributions of spatial and transcriptomic information, we observed that as the spatial scale increased, clustering results showed a gradual increase in spatial neighborhood homogeneity, accompanied by a decrease in transcriptomic neighborhood homogeneity, with the former eventually surpassing the latter. SpatialZoomer also supports diverse downstream applications, such as identifying biologically meaningful signals, spatially dependent cell subclusters, and uncovering complex tissue architectures (Fig. 1b-d).

### SpatialZoomer reveals the emergence of Alzheimer’s Disease progression signal at specific scale

SpatialZoomer is capable of identifying biologically meaningful signals at specific scales, such as those related to disease progression in perturbation or time-series scenarios. Here, we applied SpatialZoomer to an Alzheimer’s disease (AD) progression model consisting of six mouse brain samples profiled using the Xenium v1 platform with a 347-gene panel. The dataset includes both wild-type and TgCRND8 Alzheimer’s disease mouse models collected at three time points, representing mild, moderate, and advanced stages of amyloid-beta (Aβ) deposition.

We demonstrated that SpatialZoomer could localize AD progression-related signals at specific scale (Fig. 2a). At low scales (0.0, 1.1, 3.0, and 4.5), AD progression-related signal was obscured by batch variations such as developmental differences or tissue sectioning locations. Proportions of samples within clusters were relatively uniform, failing to capture the expected enrichment of disease-related clusters in the later disease stages (Fig. 2b–c). However, at a higher scale of 6.0, a distinct and biologically significant cluster emerged, identified as the AD progression-related cluster (Fig. 2b–c). This cluster was enriched in the moderate and advanced AD stages, with limited presence in the mild-stage and wild-type controls (Fig. 2b-c). As the scale increased further (8.0, 10.5), the signal became more enriched, appearing exclusively in the moderate and advanced stages.

**Figure 2.**
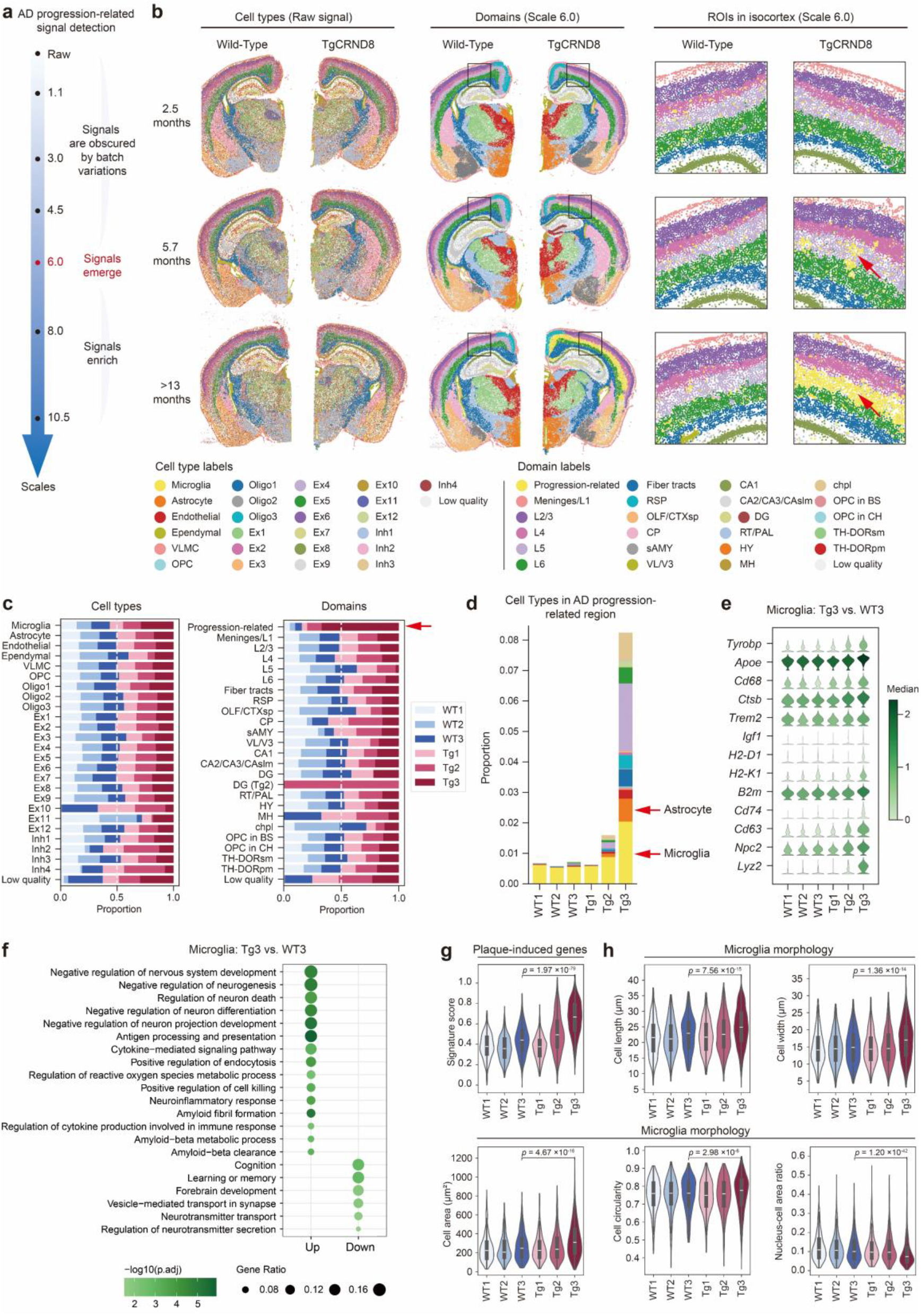
Emergence of Alzheimer’s disease progression-related signal at specific scales. **a**, Schematic illustrating the detection of AD progression-related signals across different scales. At low scales, the signals are obscured by batch variations. At scale 6.0, the signals emerge and become enriched at higher scales. **b**, Clustering results from SpatialZoomer based on raw signals (left) and signals at scale 6.0 (middle and right), representing cell type and domain scales, respectively. The right panel shows a zoomed-in view of the clustering results at scale 6.0, corresponding to the area highlighted by the black box in the middle panel. VLMC, Vascular leptomeningeal cells; OPC, oligodendrocyte progenitor cells; Oligo, oligodendrocytes; Ex, excitatory neuron; Inh, inhibitory neuron; RSP, retrosplenial area; OLF, olfactory area; CTXsp, cortical subplate; CP, Caudoputamen; sAMY, Striatum-like amygdalar nuclei; VL, lateral ventricle; V3, third ventricle; CAslm, CA stratum lacunosum-moleculare; DG, dentate gyrus; RT, reticular nucleus of the thalamus; PAL, pallidum; HY, hypothalamus; MH, medial habenula; chpl, choroid plexus; BS, brain stem; CH, cerebrum; TH-DORsm, thalamus, sensory-motor cortex related; TH-DORpm, thalamus, polymodal association cortex related. Cells and domains were manually annotated with reference to the Allen Brain Atlases^20^. **c**, Proportions of samples within clusters for raw signals and signals at scale 6.0. WT1, WT2, and WT3 represent wild-type samples, while Tg1, Tg2, and Tg3 represent TgCRND8 AD samples, with 1, 2, and 3 corresponding to three time points representing mild, moderate, and advanced stages. **d**, Bar plot showing the proportions of cell types in six samples within the AD progression-related cluster. **e**, Violin plot showing upregulated genes in microglia from the Tg3 sample compared to WT3 sample in the AD progression-related cluster. **f**, Enriched Gene Ontology (GO) terms for up- or downregulated genes in microglia from the Tg3 sample compared to WT3 sample in the AD progression-related cluster. **g**, Violin plot showing the signature score of public plaque-induced genes in microglia of the AD progression-related cluster across six samples. **h**, Violin plot depicting microglia morphology in the AD progression-related cluster across six samples. *P* values in panels **g**-**f** were calculated using the two-sided Wilcoxon Rank-Sum test between Tg3 and WT3.

At scale 6.0, the AD-progression related cluster was primarily localized in the L5/L6 region of the isocortex, with small fractions in the hippocampus and cortex subplate in the advanced stage^20^ (Fig. 2b). In the moderate stage, these cells formed patch-like clusters within the L5/L6 region. This cluster consisted primarily of microglia, astrocytes, and excitatory/inhibitory neurons, with microglia and astrocytes previously recognized as key contributors to AD progression^21,22^ (Fig. 2d). Differential gene expression analysis between advanced AD and corresponding wild-type samples revealed that microglia in the AD progression-related cluster were in activated state, marked by upregulation of *Tyrobp, Apoe*, and *Cd68*^21,22^ (Fig. 2e). This activation was accompanied by enhanced antigen processing and presentation (*H2-D1, H2-K1, B2m*, and *Cd74*), neuroinflammatory response (*Tyrobp, Trem2*, and *Igf1*), endocytosis activity and amyloid-beta clearance (*Apoe, Cd63, Npc2*, and *Lyz2*) (Fig. 2e-f). However, cognition, learning or memory and synaptic processes, including neurotransmitter secretion and transport, were significantly downregulated (Fig. 2f). Using publicly plaque-induced genes for scoring, we observed a significantly stepwise increase in expression levels from mild to advanced AD stages^23^ (Fig. 2g). Single-cell morphological analysis confirmed that microglia in advanced AD exhibited increased cell size, with larger length, width and area, a more rounded shape, and a reduced nucleus-to-cell area ratio (Fig. 2h).

We tested Points2Regions, a multi-scale analysis method that integrates gene expressions across bins of varying sizes^18^, on the AD dataset for comparison with SpatialZoomer. After excluding clusters associated with batch effects, Points2Regions failed to detect signals strongly related to AD progression at non-single-cell scales (5, 10, 15, and 20). Under the same number of scales and clustering resolutions, SpatialZoomer completed the analysis for 3 clustering resolutions and 11 scales in 31.66 minutes with a peak memory usage of 17.56 GB, while Points2Regions with the optimization strategy required 62.14 minutes and 38.10 GB. Collectively, these results highlight SpatialZoomer’s ability to detect scale-specific signals in this scenario, superior computational efficiency, and low memory consumption.

### SpatialZoomer detects spatially dependent subclusters at single-cell scale

As the spatial scale increases, SpatialZoomer progressively integrates spatial contexts from broader neighborhoods into each cell’s transcriptomic profile, potentially enabling cells from a single cluster at scale 0 to separate into spatially dependent subclusters at larger scales. These spatial contexts can amplify transcriptomic heterogeneity between biologically distinct subtypes.

In a Xenium lung cancer dataset with 377 genes, SpatialZoomer identified macrophages as spatially dependent clusters that separated into alveolar (AM) and interstitial macrophage (IM) subtypes. For each cluster identified in the raw signal space, we calculated the variance of cell distributions across UMAP embeddings at different scales (Fig. 3a-b). Spatial dependency was then quantified by fitting slopes of the variance–scale curves, which enabled ranking of clusters (Fig. 3b). Macrophages, the top-ranked cluster, appeared as a single cluster at small scales but separated into two distinct subclusters as the scale increased (Fig. 3c). The elbow method identified scale 4.5 as the critical separation point, and density peak clustering identified two subclusters, C0 and C1, at this scale (Fig. 3d–e). A Ward’s linkage-based hypothesis test confirmed that the separation was statistically significant compared with the raw signal (*P* = 0.00; Fig. 3d).

**Figure 3.**
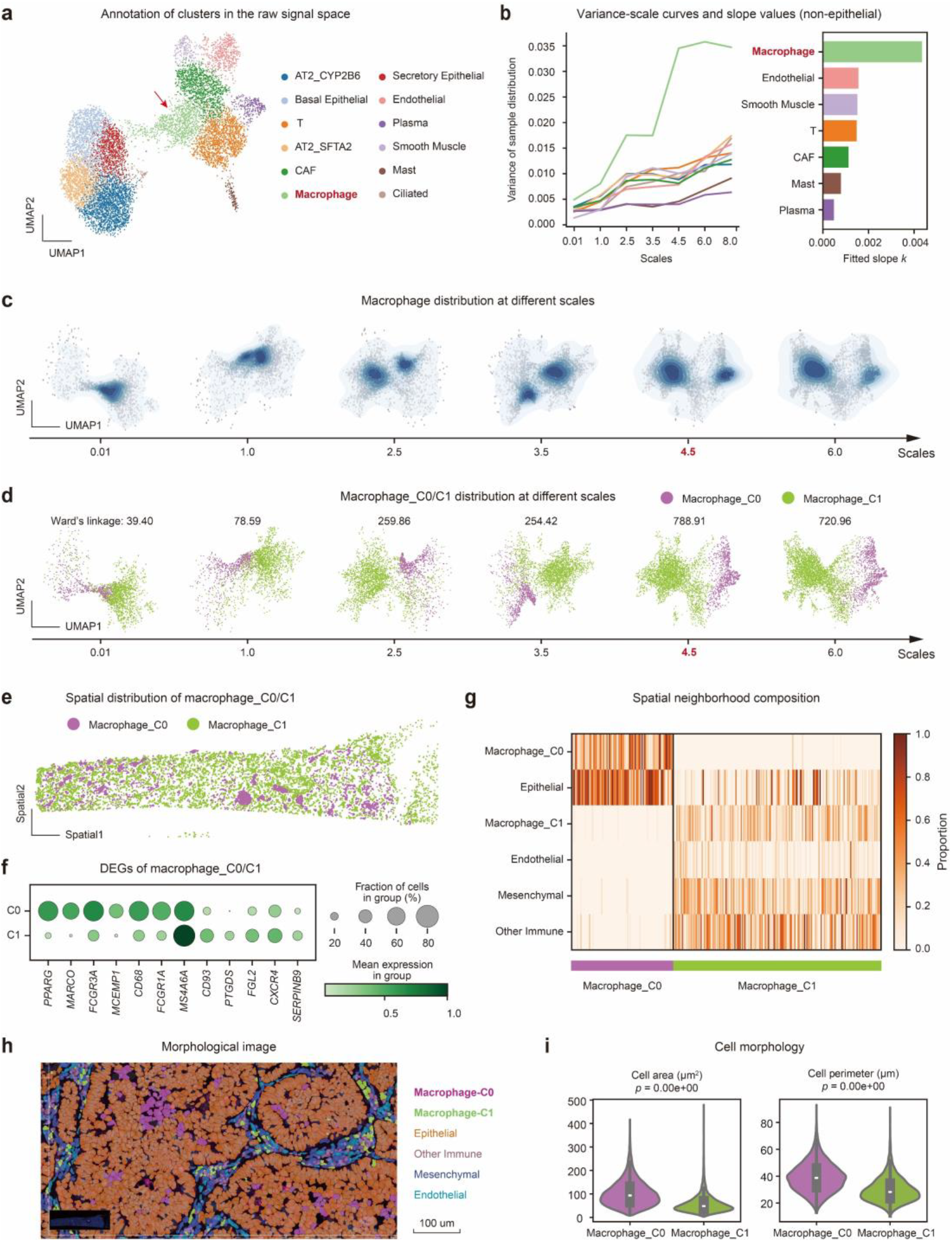
Identification of spatially dependent macrophage subclusters in lung cancer. **a**, Clusters identified by SpatialZoomer in the raw signal space. Cell types were manually annotated using marker genes. **b**, Variance-scale curves for non-epithelial cell clusters, with fitted slopes quantifying the potential for cluster separation as spatial scale increases. **c**, UMAP with kernel density estimation (KDE) showing macrophage distributions across scales. **d**, UMAP showing the distribution of two macrophage subclusters, Macrophage_C0 and Macrophage_C1, across scales. The subclusters were identified at the critical separation scale of 4.5. Ward’s linkage values were labeled, with higher values indicating stronger separation. **e**, Spatial distribution of macrophage subtypes. **f**, Differentially expressed genes between macrophage subtypes (logFC > 0.2, adjusted *P* value < 0.05). To minimize background noise, only genes with higher mean expression in macrophages compared to other cell types were included. **g**, Spatial neighborhood composition based on the 10 nearest neighbors per cell. **h**, Morphological image with cell type annotations. DAPI and cell boundary staining with cell type-specific coloring by Xenium Explorer highlight morphological and neighborhood differences between the two macrophage subtypes. **i**, Violin plots showing cell area and perimeter for the two subtypes. *P* values were assessed using Welch’s t-test.

Multiple lines of evidence indicated that C0 and C1 correspond to biologically distinct AM and IM subtypes. (1) For transcriptomics, C0 highly expressed canonical AM marker *PPARG* and *MARCO*, consistent with functions in phagocytosis of inhaled pathogens/particles and immune regulation^24,25^ (Fig. 3f). By contrast, C1 expressed *CD93, CXCR4*, and *PTGDS*, genes linked to chemotaxis, tissue repair, and immune responses, consistent with IM features. Scoring the human lung single-cell atlas with the differentially expressed gene sets further supported these subtype-specific signatures^26^. (2) For spatial distribution, C0 and C1 exhibited distinct aggregation patterns (Fig. 3e). Neighborhood composition revealed that C0 localized within alveolar spaces adjacent to epithelial cells, whereas C1 localized in interstitial regions enriched with endothelial, mesenchymal, and immune cells, consistent with AM and IM positioning (Fig. 3g–h). (3) For morphology, C0 exhibited significantly larger cell area and perimeter than C1 (Fig. 3h–i). Because the probe panel in this dataset covered only hundreds of genes, direct identification of fine-grained subtypes was challenging. By integrating spatial context along the scale axis, SpatialZoomer overcame this limitation and enhanced cell state characterization.

In a Xenium ovarian cancer dataset comprising 5,101 genes, which provides a considerably broader panel compared with the lung cancer dataset analyzed above, SpatialZoomer identified CAF as a spatially dependent cluster (Ovary_S1, ovarian papillary serous carcinoma). As the spatial scale increased, CAF subclusters progressively separated in feature space, and two distinct subtypes, CAF_C0 and CAF_C1, were identified at the critical separation scale of 6.5 (Fig. 4a-b). These subtypes exhibited heterogeneity across spatial distributions, morphology, and transcriptomic profiles. Spatial distribution revealed that CAF_C0 predominantly localized adjacent to SOX2-OT^+^ tumor cells and was embedded within extracellular matrix (ECM) regions exhibiting lighter hematoxylin and eosin (H&E) staining (Fig. 4c-e). By contrast, CAF_C1 colocalized with pericytes within stromal gaps among tumor nests comprising tumor cells and proliferating tumor cells (Fig. 4c-e). For morphology, CAF_C1 demonstrated larger cell and nuclear areas and a more regular shape than CAF_C0 (Fig. 4f). Differential gene enrichment analysis revealed that CAF_C0 upregulated genes associated with ECM organization, while CAF_C1 upregulated genes associated with responses to external stimuli, hypoxia, and nutrient levels (Fig. 4g). To validate these findings, we analyzed an independent ovarian cancer dataset (Ovary_S2, ovarian adenocarcinoma). SpatialZoomer again identified two spatially dependent CAF subclusters (Fig. 4h). Scoring CAFs in Ovary_S2 with CAF_C0 and CAF_C1 signature genes from Ovary_S1 revealed highly consistent spatial distribution patterns, matching the two subtypes across datasets (Fig. 4i–j). Together, even in datasets with broad gene coverage, SpatialZoomer’s multi-scale framework effectively leverages spatial contexts to amplify molecular signals, enabling the identification of spatially and functionally distinct subpopulations.

**Figure 4.**
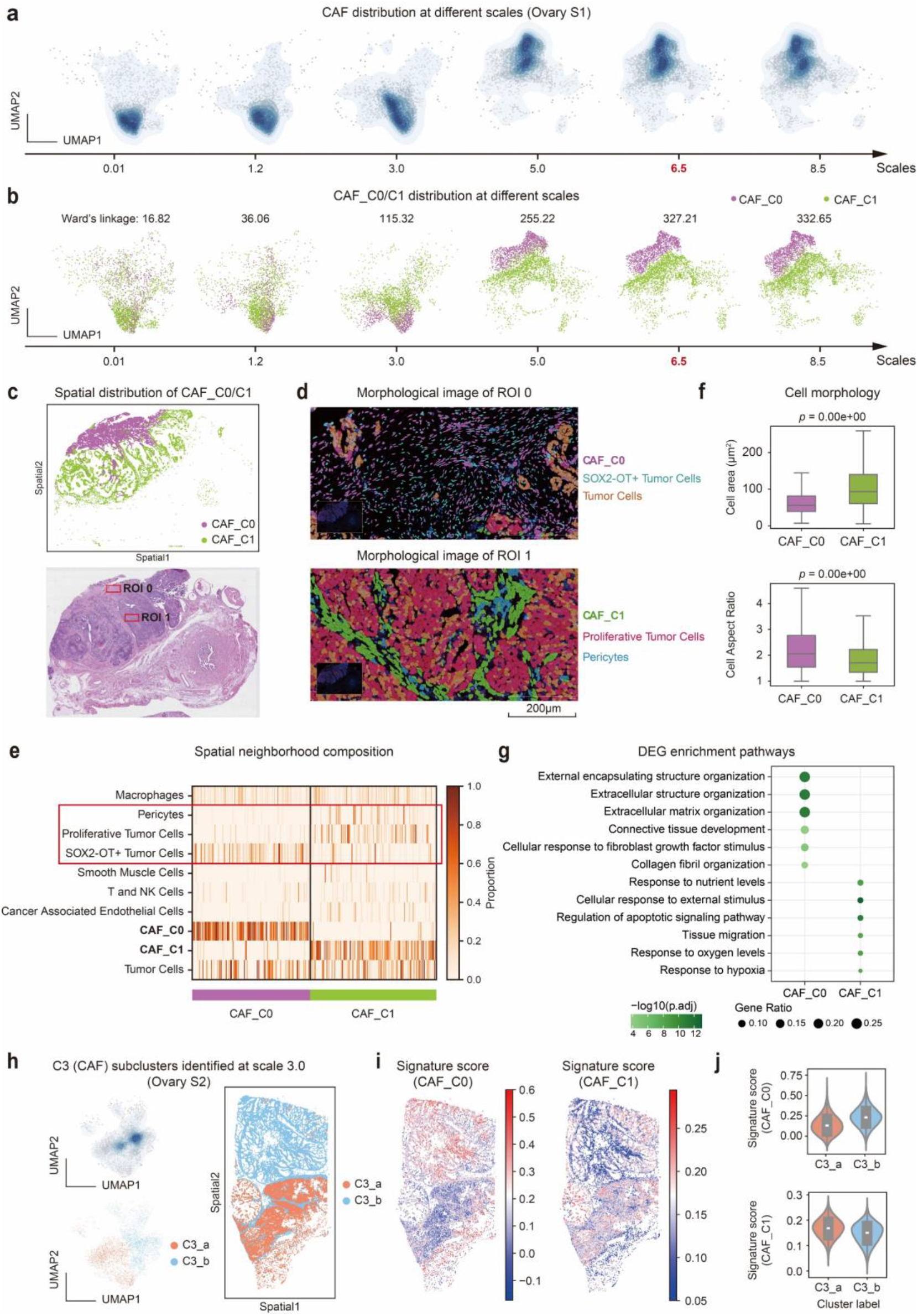
Identification of spatially dependent CAF subclusters in ovarian cancer. **a**, UMAP with KDE showing CAF distributions across scales in sample Ovary S1. **b**, UMAP showing the distribution of two CAF subclusters, CAF_C0 and CAF_C1, across scales. The subclusters were identified at the critical separation scale of 6.5. Ward’s linkage values were labeled, with higher values indicating stronger separation. **c**, Spatial distribution of CAF subtypes and the H&E-stained image. Representative regions of CAF_C0 and CAF_C1 are marked as ROI 0 and ROI 1. ROI, Regions of Interest. **d**, Morphological images of ROI 0 and ROI 1. DAPI and cell boundary staining with main cell type-specific coloring by Xenium Explorer show the typical neighborhood of CAF_C0 and CAF_C1. **e**, Spatial neighborhood composition based on the 10 nearest neighbors per cell. **f**, Boxplots showing nucleus areas and cell densities for the two subtypes. *P* values were assessed using Welch’s t-test. **g**, GO enrichment analysis of differentially expressed genes (DEGs) between CAF_C0 and CAF_C1 (logFC > 0.2, adjusted *P* value < 0.05). To minimize background noise, only genes with higher mean expression in CAFs compared to other cell types were included. Pathways with adjusted *P* value < 0.01 were considered significant. **h**, Subclusters of C3 (CAF), designated as C3_a and C3_b, identified in sample Ovary S2 at the critical separation scale of 3.0. Left: UMAP showing the distributions of the cluster and its subclusters at scale 3.0. Right: spatial distribution of the two subclusters. **i**, Signature scores of CAF_C0 and CAF_C1 from Ovary S1 applied to C3 (CAF) in Ovary S2. Signatures were defined by DEGs of CAF_C0 and CAF_C1, filtered using the same thresholds as described in **g. j**, Violin plots illustrating the CAF_C0 and CAF_C1 signature scores in C3_a and C3_b subclusters.

### SpatialZoomer uncovers stress-proliferation-EMT tri-layer organization of tumor cells at the niche level

Understanding how individual cells organize into functional collectives is essential for deciphering biological systems, particularly in tumors where such organization is often highly complex. The multi-scale spatial features extracted by SpatialZoomer could be used to uncover such tissue architectures. To demonstrate this capability, we applied SpatialZoomer to a cervical cancer sample profiled using the Xenium Prime platform with the 5K-gene panel. At the niche level (scale 6.0), SpatialZoomer revealed multilayered luminal structures, with three representative ROIs selected for detailed visualization (Fig. 5a–b). These luminal structures exhibited a conserved organization: stromal and immune cells occupied the core, surrounded by tumor cells stratified into stressed, proliferative, and EMT-like subtypes arranged radially from center to periphery, each defined by distinct gene expression programs (Fig. 5c–d).

**Figure 5.**
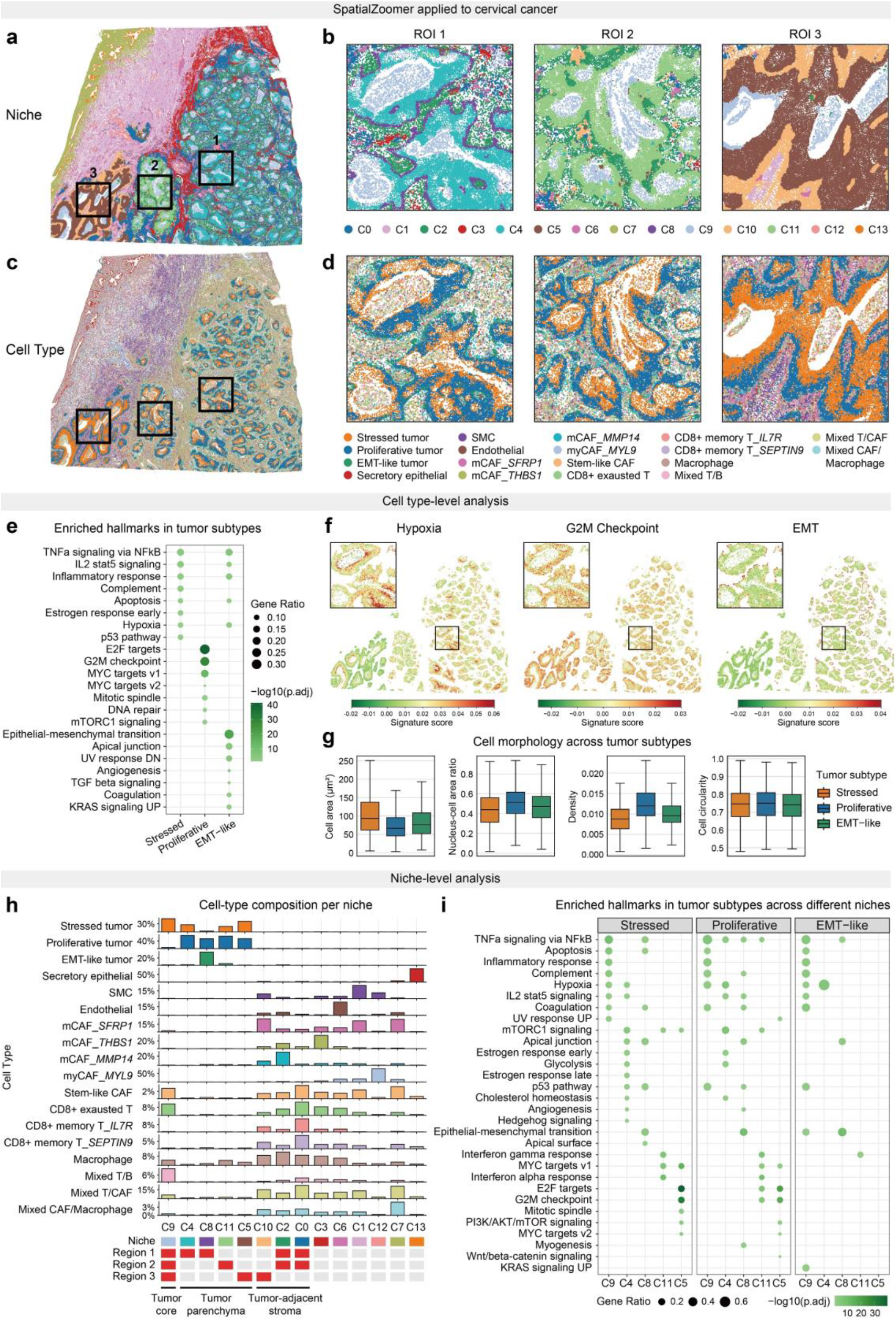
Identification of the stress-proliferation-EMT tri-layer organization formed by tumor cells at the niche level. **a-b**, Niche-scale results from SpatialZoomer at scale 6.0. Subpanel b presents zoomed-in views of three ROIs highlighted in a. **c-d**, Cell-scale results from SpatialZoomer using raw signals, with d showing zoomed-in views of ROIs corresponding to c. mCAF, matrix CAF; myCAF, myofibroblastic CAF. **e**, Enriched hallmarks resulting from upregulated genes in each tumor subtype compared to the other two. **f**, Spatial scatter plot displaying signature scores for hypoxia, G2M checkpoint, and EMT across single cells. The black inset in the upper left corner presents the magnified ROI 1. **g**, Boxplots showing differences in cell morphology across the three tumor subtypes, including metrics such as cell area, nucleus-to-cell area ratio, cell density, and circularity. For each cell morphology descriptor, pairwise comparisons among the three tumor subtypes revealed statistically significant differences (*P* = 0.00, Welch’s t-test). **h**, Bar plots showing the composition of cell types across 14 niches. Values are column-normalized, with colors corresponding to cell types in c-d. The bottom heatmap summarizes niche information, with the first row indicating niche colors in a-b. Rows 2-4 classify niches based on their spatial location within three regions corresponding to ROI 1 to ROI 3, while columns represent niches arranged sequentially as tumor core, tumor parenchyma, and tumor-adjacent stroma. Region 1: C9-C4-C8-C2-C0, Region 2: C9-C11-C2-C0, and Region 3: C9-C5-C10. **i**, Enriched hallmarks resulting from upregulated genes compared across different niches within the same tumor cell subtype.

Stressed tumor cells were characterized by G0/G1 arrest, upregulation of *CDKN1A*, and activation of stress-associated pathways, including hypoxia, apoptosis, and p53 signaling (Fig. 5e-f). Proliferative tumor cells activated cell cycle programs, including G2/M checkpoint, E2F targets, MYC targets, and DNA repair hallmarks. EMT-like tumor cells showed upregulation of EMT and TGF-β signaling (Fig. 5e-f). Beyond their transcriptional differences, the three tumor cell subtypes also exhibited different morphological features. Stressed tumor cells displayed larger cell areas, likely due to intracellular material accumulation and expanded organelle content under p53 pathway activation and hypoxic stress (Fig. 5g). In contrast, proliferative tumor cells appeared smaller, with increased nucleus-to-cell ratios, reduced nuclear solidity, and dense local packing, consistent with active cell cycling (Fig. 5g). Meanwhile, EMT-like tumor cells lost morphological regularity, exhibiting reduced cell circularity, solidity, and symmetry, in line with the polarity loss, cytoskeletal remodeling, reduced adhesion, and motility activation characteristic of EMT (Fig. 5g).

Tumor-associated niches exhibited spatial heterogeneity across the three regions corresponding to the selected ROIs (Fig. 5a-b, h). While all regions contained luminal structure-associated niches composed of stromal cells and three tumor subtypes, only the tumor core niche C9 was consistently shared across regions (Fig. 5a-b, h). By contrast, the surrounding tumor parenchyma displayed region-specific niche compositions: C4 and C8 in region 1, C11 in region 2, and C5 in region 3 (Fig. 5h). The coexistence of a conserved core alongside spatially distinct peripheral niches suggests that niche heterogeneity may be shaped by variations in the adjacent stroma or broader tissue context (Fig. 5h).

To assess whether these spatially distinct niches exhibit biological variations, we compared niches within the same tumor cell subtype (Fig. 5i). Interestingly, although stressed, proliferative, and EMT-like tumor cells represent distinct transcriptional states, they exhibited consistent and niche-specific gene expression profiles when located within the same niche (Fig. 5i). For example, in region 1, all tumor cell subtypes within niche C4 were consistently enriched for hypoxia, IL2–STAT5 signaling, and glycolysis, whereas those in niche C8 displayed elevated EMT activity (Fig. 5i). In region 2, niche C11 showed upregulation of interferon α/γ responses and MYC target V1 genes, whereas in region 3, niche C5 showed activated cell cycle activity. Moreover, we observed a gradual and global increase in cell cycle activity across niches from region 1 to 3, transitioning from C4 to C11 to C5. Together, these findings suggest that tumor cells form an organized stress-proliferation-EMT tri-layer structure, and that spatially distinct niches with similar cellular compositions can exhibit distinct transcriptomic profiles likely shaped by their spatial contexts.

## ·Discussion

SpatialZoomer is a computational framework for multi-scale spatial analysis of single-cell resolution spatial transcriptomics data, introducing spatial scale as an analytical axis to uncover diverse biological insights. SpatialZoomer naturally integrates spatial information and high-dimensional transcriptomic data by modeling gene expressions as signals on a spatial neighbor graph. It applies heat kernel-based low-pass filters to extract spatial features at different scales. To select critical scales in a data-driven manner, SpatialZoomer employs dynamic programming to partition candidate scales into intervals and selects critical scales from each. Much like adjusting the focus of a microscope, SpatialZoomer enables the identification of cell types, niches, and domains at different scales, as well as the detection of patterns such as disease progression-related signals that emerge at specific scales. Moreover, by scanning along the scale axis, SpatialZoomer enhances the raw signals by incorporating spatial contexts and reveals spatially dependent cell subtypes, particularly within complex tumor microenvironment. The systematic nature of this multi-scale perspective further enables the characterization of complex tissue architecture.

SpatialZoomer enhances the resolution of cell subtype identification by incorporating spatial context at multiple scales. The identified spatially dependent subclusters are not merely localized in different microenvironments or tissue domains, but also exhibit distinct transcriptomic profiles and morphological characteristics, highlighting that spatial contexts play a key role in shaping cellular heterogeneity. The importance of spatial contexts in refining cell states has also been emphasized by BANKSY^11^, a method that augments each cell’s gene expression profile with the averaged neighborhood expression and spatial gradient contexts. Unlike BANKSY, which relies on feature fusion from three fixed spatial scales, SpatialZoomer explicitly introduces a spatial scale axis. By tracking how cellular features vary as progressively larger spatial contexts are incorporated, SpatialZoomer reveals whether and when subclusters emerge. Notably, SpatialZoomer demonstrates robust performance across diverse spatial scenarios, adapting to different patterns of spatial organization. It successfully detects macrophage subclusters in lung cancer due to niche-driven separation and CAF subclusters in ovarian cancer due to domain-driven separation.

SpatialZoomer also demonstrates the ability to uncover cellular spatial organization. In cervical cancer, it identified a stress-proliferation-EMT tri-layer structure, along with regions showing a simpler stress–proliferation bi-layer structure, the latter of which was also observed in a lung cancer sample. A similar spatial organization was previously reported in oral squamous cell carcinomas (OSCC) using 10x Visium ST data^27^, described as tumor core, transitory zone, and leading edge. However, because Visium captures spot-level signals with mixed cells, proliferative and EMT programs were both interpreted as features of the leading edge. Our single-cell resolution analysis revealed that the EMT layer in cervical cancer is thin and spatially distinct from the proliferative layer, providing a refined view of tumor architecture and potentially complementing previous interpretations. Interestingly, both datasets used in this study and those previously reported are derived from squamous cell carcinomas (SCC) originating from different organs, and the tumor cells in both exhibit basal-like transcriptomic features^27^. These observations suggest that this spatial organization pattern may be a recurring feature of SCC, potentially pointing to shared biological mechanisms and therapeutic vulnerabilities.

SpatialZoomer can also be applied to sequencing-based ST platforms such as Visium HD and Stereo-seq. However, because spot-level data contain mixed signals, intermediate states are commonly observed at different scales. Addressing this challenge may require tailored modeling of mixed signals before applying the multi-scale framework. In addition, this work incorporated single-cell-level morphological information, including shape descriptors derived from cell and nuclear segmentation, and local cell density based on spatial coordinates. These features served as strong supporting evidence for validating cellular heterogeneity. Given that spatial transcriptomics (ST) data are often paired with H&E-stained histological images, and with the rapid emergence of multimodal spatial omics technologies^28,29^, the potential of SpatialZoomer in integrative multi-omics analysis remains to be explored.

In summary, SpatialZoomer offers an accurate, versatile, scalable, and computationally efficient framework for multi-scale analysis of single-cell ST data.

## Data availability

The CosMx SMI mouse coronal hemisphere sample was downloaded from https://nanostring.com/products/cosmx-spatial-molecular-imager/ffpe-dataset/cosmx-smi-mouse-brain-ffpe-dataset/. The vizgen MERFISH mouse brain dataet was downloaded from https://info.vizgen.com/mouse-brain-map. The Xenium Coronal whole brain with a 248-gene panel was downloaded from https://www.10xgenomics.com/datasets/fresh-frozen-mouse-brain-replicates-1-standard. The Xenium Prime 5K coronal hemisphere sample was downloaded from https://www.10xgenomics.com/datasets/xenium-prime-fresh-frozen-mouse-brain. The Xenium Alzheimer’s Disease Mouse Model dataset, consisting of six coronal hemispheres and profiled with a 347-gene panel, was downloaded from https://www.10xgenomics.com/datasets/xenium-in-situ-analysis-of-alzheimers-disease-mouse-model-brain-coronal-sections-from-one-hemisphere-over-a-time-course-1-standard. The Xenium lung cancer sample with a 377-gene panel was downloaded from https://www.10xgenomics.com/datasets/ffpe-human-lung-cancer-data-with-human-immuno-oncology-profiling-panel-and-custom-add-on-1-standard. The human lung single cell atlas (LungMAP)^26^ was downloaded from https://www.lungmap.net/omics/?experiment_id=LMEX0000004396. Two Xenium Prime 5K ovarian samples were downloaded from https://www.10xgenomics.com/datasets/xenium-prime-ffpe-human-ovarian-cancer and https://www.10xgenomics.com/datasets/xenium-prime-fresh-frozen-human-ovary. The Xenium Prime 5K cervical cancer dataset was downloaded from https://www.10xgenomics.com/datasets/xenium-prime-ffpe-human-cervical-cancer. The Xenium non-small cell lung cancer sample with a 480-gene panel was downloaded from https://www.10xgenomics.com/datasets/ffpe-human-lung-cancer-data-with-human-immuno-oncology-profiling-panel-and-custom-add-on-1-standard.

## Code availability

The SpatialZoomer Python package and tutorials are available at: http://lifeome.net/software/spatialzoomer. An interactive demo showcasing multi-scale results along a sliding scale axis can be accessed at https://li-xinqi.github.io/SpatialZoomer/.

## Acknowledgements

This work was supported by the National Key Research and Development Program of China (Nos. 2020YFA0712403, 2021YFF1200901), the National Natural Science Foundation of China (Nos. 62133006, 92268104), Beijing Natural Science Foundation (No. Z230015).

## Author contributions

J.G. conceptualized and supervised the study. X.L. and Y.F. developed the computational framework and performed bioinformatic analyses. J.G., X.L., and Y.F. discussed the methods and results, and drafted the original manuscript with the assistance of other authors. Y.H. and J.H. contributed to algorithm development. W.G., N.Y., Z.C. (Tsinghua university), Y.W., and Y.M. assisted in interpreting the results. C.Z. (Peking university), X.Z. and L.H. provided valuable suggestions during the study. All authors reviewed the final version of the manuscript.

## Methods

### Processing and analysis of spatial transcriptomics data

#### Data preprocessing

All data preprocessing steps were conducted using “squidpy” (version 1.5.0). Cell quality control retained cells with total genes and transcript counts above predefined thresholds and removed those with a “nucleus_area” value of zero in Xenium dataset. Gene quality control filtered out lowly expressed genes based on the number of cells in which they were detected. Following quality control, the raw count matrix was normalized to the median count depth across cells and subsequently subjected to log transformation.

#### Differential gene expression analysis

Differentially expressed genes (DEGs) were identified using the “rank_genes_groups” function in Scanpy^30^ (version 1.10.1) with an adjusted *P* value < 0.05. The *P* values were determined using the Wilcoxon rank-sum test and adjusted by the Benjamini-Hochberg (BH) correction. Up- or down-regulated genes were subsequently input into pathway enrichment analysis using the “enricher” function from the clusterProfiler R package^31^ (version 4.3.2). Enriched terms were identified as significant with an adjusted *P* value < 0.05, determined by a one-sided Fisher’s exact test and adjusted using the BH correction. The candidate functional terms were derived from Hallmark and Gene Ontology (GO) terms in the MSigDB database^32^.

#### Signature score calculation

Signature scores were calculated using the “tl.score_genes” function from the Scanpy package^30^. In the ovarian cancer analysis (Fig. 4), gene signatures were derived from DEGs between CAF_C0 and CAF_C1 subclusters, with thresholds set at an adjusted *P* value < 0.05 and logFC > 0.2. To minimize background noise, particularly from genes introduced by inappropriate cell segmentation that may capture expression from neighboring cells, only genes with higher mean expression in CAFs compared to all other cell types were retained. In the cervical cancer analysis, gene sets were obtained from Hallmark gene sets in the MSigDB database^32^, intersected with the Xenium 5K gene panel.

#### Additional data processing

For the Alzheimer’s disease dataset, spatial coordinates from six mouse brain samples were concatenated into a common coordinate space by rotation and translation, allowing SpatialZoomer to be applied jointly across all samples.

For cell type annotation, initial labeling for the mouse brain, lung cancer, and cervical cancer datasets was performed using CellTypist^33^. Annotations were then manually verified and refined based on cluster-specific differentially expressed genes and the expression of known marker genes. For the ovarian cancer sample (S1), cell type annotations were obtained from the official dataset release provided by 10x Genomics (https://www.10xgenomics.com/datasets/xenium-prime-ffpe-human-ovarian-cancer).

